# Hyperface: a naturalistic fMRI dataset for investigating human face processing

**DOI:** 10.64898/2026.03.11.711073

**Authors:** Matteo Visconti di Oleggio Castello, Guo Jiahui, Ma Feilong, Manon de Villemejane, James V. Haxby, Maria Ida Gobbini

## Abstract

Faces convey information that guides social behavior, yet neuroimaging studies investigating human face processing typically use static images with small sets of identities under artificial conditions. Controlled designs limit our ability to characterize human face processing under naturalistic conditions or test whether computational models generalize beyond the laboratory. To address this gap, here we release *hyperface*, a naturalistic face viewing fMRI dataset designed to investigate human face processing in response to faces portrayed in videos mimicking more ecologically valid conditions. Twenty-one participants watched 707 unique face video clips that vary systematically in identity, gender, age, ethnicity, expression, and head orientation. Each clip was rated by independent observers, and pairwise similarity judgments were collected through a behavioral arrangement task. Technical validation confirms high data quality with low motion, high tSNR, and high inter-subject correlation in visual and face-processing regions. The *hyperface* dataset is part of a comprehensive experimental framework to investigate human face processing: all 21 participants also watched “The Grand Budapest Hotel,” performed a dynamic face localizer task, and 10 participants completed an additional face identity task with personally familiar and visually familiarized faces. These datasets are publicly available and enable within-subject comparisons across paradigms. Together they provide a unique resource for characterizing human face processing under naturalistic conditions and for benchmarking computational models against human brain responses.

## Background & Summary

Faces convey rich information that guides our behavior in social contexts. From a single face we can quickly recognize identity, and infer gender, age, emotional state, personality traits, and other socially relevant characteristics of an individual.^1–5^ Few other visual objects can convey such rich and socially relevant information. To understand the functional representations underlying face processing, neuroimaging studies typically use small sets of static images of identities presented under highly controlled conditions.^6,7^ This controlled approach has been highly productive, and it has allowed researchers to map the distributed face-processing network and its functional representation of face identity, expression, and personal familiarity.^3,8–11^ However, controlled experimental designs limit our ability to fully characterize face processing under dynamic, naturalistic conditions, or to test whether computational models of face processing generalize to real-world scenarios.

Naturalistic neuroimaging paradigms address these limitations by enabling investigation of brain function under conditions that better resemble real life.^12–15^ However, only a few publicly available datasets so far exist to investigate face processing under naturalistic conditions.^16,17^ To address this gap, here we release a naturalistic face viewing fMRI dataset called *hyperface*, in which 21 participants watched 707 unique 4 s face video clips sampled from publicly available YouTube interviews. To investigate face processing under a wide range of conditions, these clips show individuals who vary systematically in identity, gender, age, ethnicity, expression, and head orientation. Each clip was rated by independent observers for perceived gender, age, ethnicity, head orientation, facial expression, and number of people present in each scene. Additional pairwise similarity judgments were collected through a behavioral arrangement task in which independent observers were asked to arrange the clips based on perceived similarity of face appearance. In recent work,^18^ we used this dataset with hyperalignment and representational similarity analysis to show that DNNs trained for face and object identification tasks successfully capture human cognitive categorical face representation in the behavioral arrangement task, but fail to explain brain representations of identity during dynamic, naturalistic face viewing. This suggests that state-of-the-art visual DNNs are not yet accurate models of human face processing under naturalistic conditions, capturing only part of the reliable functional representations evoked by dynamic faces.^18–20^

The *hyperface* dataset is part of a comprehensive experimental framework designed to characterize human face processing under naturalistic conditions.^21^ This framework includes two additional fMRI datasets already publicly available: a movie-watching dataset in which participants watched Wes Anderson’s “The Grand Budapest Hotel,”^17^ a socially rich movie with many characters and social relationships; and a controlled fMRI paradigm in which participants viewed faces of personally familiar and visually familiarized individuals with different head orientations.^9^ Importantly, all 21 *hyperface* participants also watched “The Grand Budapest Hotel”, performed a dynamic face localizer task, and 10 participants completed all experiments.^21^ This comprehensive set of experiments enables testing hypotheses about face processing across paradigms, and provides a unique data resource across human neuroscience, computational neuroscience, and artificial intelligence. It allows investigating human face processing with a variety of methods, such as inter-subject correlation (ISC),^22^ multivoxel pattern analysis and representational similarity analysis,^23,24^ or voxelwise encoding models;^15,25,26^ and it enables benchmarking computational models of face processing against human brain responses and behavioral data.^18^ This Data Descriptor provides the first documented release of the *hyperface* data; the previously released datasets used different stimuli and do not include the face-clip data released here.

## Methods

### Participants

Twenty-one participants (11 women, mean age 27.3 ± 2.3 years, range 22–31) took part in the experiment. All had normal or corrected-to-normal vision and used a custom-fitted CaseForge headcase (https://github.com/gallantlab/headcase-pipeline) to minimize head motion. The study was approved by the Dartmouth Committee for the Protection of Human Subjects under protocol STUDY00029780. All participants provided written informed consent to participate and to share their de-identified data. The consent form stated that data may be shared with other researchers or deposited in a publicly available data archive with personal identifiers removed.

The same 21 participants also watched “The Grand Budapest Hotel” in a separate scanning session (see ref.^17^ for more information), and 10 of the 21 participants performed a face identity task with personally familiar and visually familiarized faces in two separate scanning sessions (see ref.^9^ for more information). These additional sessions used different stimuli than the naturalistic clips used here, and the tasks were not based on performance. Therefore, participation in these additional sessions is unlikely to bias brain responses in the present dataset.

### Stimuli

We created 707 unique face video clips (4 s each, no audio) by sampling publicly available YouTube interviews. The 707 clips were curated in two passes to assemble a large and diverse set of naturalistic faces spanning two fMRI sessions. Thus, this number of stimuli reflects the selection process rather than a predetermined target. The clips showed different individuals varying in age, ethnicity, gender, expression, and head orientation. Clips were cropped as necessary to remove distracting text, and audio was removed. Because these clips are short excerpts of YouTube videos, the video files are not redistributed with this data release. Instead, the full provenance of every clip is provided together with code to regenerate and verify the clips from their original sources (see Data Records).

#### Behavioral rating task

A total of 121 Amazon Mechanical Turk workers participated in a behavioral rating task. In each trial, workers watched a video clip and rated the stimulus on six features: perceived gender (male/female), age (0–10, 11– 20, 21–30, 31–40, 41–50, 51–60, 61–70, 70+), ethnicity (White, Black or African American, Asian, Indian, Hispanic or Latino, Other), expression (neutral, happiness, surprise, anger, disgust, sadness, fear), number of people (1, 2, more than 2), and head orientation (mostly left, mostly center, mostly right); see Figure 1B. All 707 clips were divided into 15 independent sessions (approximately 47 clips each). Each worker completed one session. At least eight workers completed each session, and the final rating for each clip was assigned as the modal response across workers. Before rating, workers answered comprehension questions, and those who failed were excluded. Inter-rater agreement (the proportion of raters matching the modal label) was high for the more objective attributes and moderate for the more subjective ones: perceived gender 0.99, ethnicity 0.90, head orientation and number of people 0.75, age 0.71, expression 0.70). Note that the provided labels are intended as coarse descriptors, and users may want to extend these annotations with more precise descriptions. For example, perceived gender was recorded as a binary judgment, but this could be extended to include non-binary options. Because facial expressions are dynamic and can be brief,^27–29^ a single categorical label may under-represent transient expressions, and researchers may choose to measure time-resolved expression ratings.

**Figure 1.**
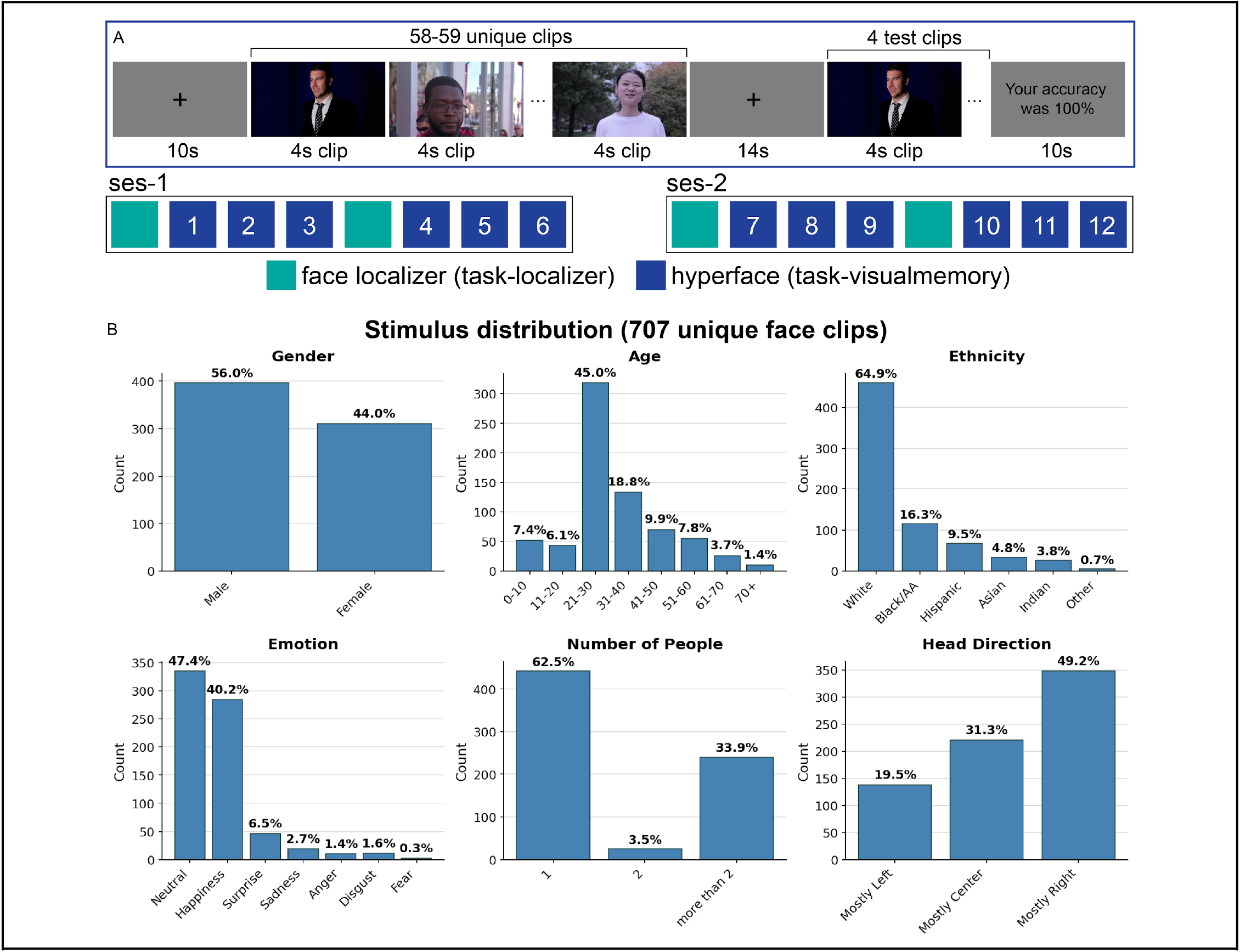
Experimental design and stimulus distribution of face stimuli. **A**. In each *hyperface* functional run participants watched 58-59 short face clips (4 s each, no sound), then performed an attention check by indicating whether four face clips had been presented in the run. Data were collected across two scanning sessions. Each session included six *hyperface* runs (*task-visual memory*) and two dynamic face localizer runs (*task-localizer*). **B**. 707 unique face clips were selected from publicly available YouTube interviews and rated on Amazon Mechanical Turk for Gender, Age, Ethnicity, Emotion, Number of People, and Head Direction. Each clip was rated by at least eight Mechanical Turk workers, and the final rating was assigned as the modal response (see Methods).

#### Behavioral arrangement task

An independent group of 39 Amazon Mechanical Turk workers performed an arrangement task to obtain pairwise similarity judgments for all face clips.^18,30^ Within each trial, all stimuli from one scanning run (58– 59 clips) were displayed as thumbnails around the circumference of a white circle on a gray background. Workers arranged the thumbnails within the circle based on perceived similarity of face appearance. Hovering over a thumbnail triggered a larger, dynamic display of the video clip; right-clicking replayed the video. Each worker completed three randomly selected trials from the 12 total runs. At least 10 workers arranged the stimuli of each scanning run. Pairwise dissimilarities were computed as on-screen Euclidean distances and averaged across workers within each run. Each worker thus produced a single, complete arrangement of all clips in a run.

### Experimental Procedure

Participants completed two scanning sessions. Each session comprised one gradient-recalled echo (GRE) fieldmap scan, eight functional runs, and one diffusion-weighted imaging (DWI) scan preceded by two spin-echo sequences with opposite phase-encoding directions for distortion correction. The first session also included one T2-weighted anatomical scan. A T1-weighted anatomical scan was collected in a separate session in which participants watched “The Grand Budapest Hotel”; this T1-weighted scan is also included in the hyperface data release. The order of the functional runs was fixed across sessions: one localizer run, three hyperface runs, one localizer run, and three hyperface runs (see Figure 1). All stimuli were back-projected onto a screen and subtended approximately 16.27° × 9.17° (W × H) of visual angle.

Following classic designs of naturalistic visual stimulation,^31–33^ participants viewed a continuous stream of at least 58 face clips (4 s each) within each hyperface run. Participants were instructed to attend to the identity of the individuals portrayed. The clips were followed by a 14-s blank period and an attention check in which four face clips were presented sequentially (see Figure 1). Participants indicated whether each individual had appeared in the preceding run. Two of the four clips were matches and two were non-matches.

Clip presentation order in the hyperface runs was counterbalanced across participants. The 707 clips were divided into 12 blocks of 58–59 clips each, one block per functional run. To ensure each block contained a balanced distribution of face attributes (gender, age, ethnicity, emotion, number of people, head direction), 10,000 random permutations of the stimulus order were generated, and we selected the permutation that minimized the summed squared deviation from a uniform distribution of attributes across blocks. Each block was also assigned four catch-trial clips: two matches (randomly selected from within the block) and two non-matches (randomly selected from a different block). The 12 blocks were then assigned to two sessions of six blocks each. Session order was counterbalanced across participants, and block order within each session was randomized for each participant.

During each localizer run, participants viewed 3-s clips of dynamically moving faces (F), bodies (B), scenes (S), objects (O), and scrambled objects (Sc).^34^ Clips were presented continuously without intervening blank periods. Participants pressed a button whenever a clip repeated. Block order followed a fixed structure: an 18-s fixation block, five category blocks (18 s each) in random order, an 18-s fixation block, and five category blocks in reversed order (e.g., B-S-F-O-Sc followed by Sc-O-F-S-B, see ref.^34^ for more details). The order of the category blocks was randomized across runs.

### Imaging Parameters

All data were acquired using a 3T Siemens Magnetom Prisma MRI scanner (Siemens, Erlangen, Germany) with a 32-channel phased-array head coil at the Dartmouth Brain Imaging Center. Functional, blood oxygenation level-dependent (BOLD) images were acquired in an interleaved fashion using gradient-echo echo-planar imaging with pre-scan normalization, fat suppression, a multiband (i.e., simultaneous multi-slice; SMS) acceleration factor of 4 (using blipped CAIPIRINHA), and no in-plane acceleration (i.e., GRAPPA acceleration factor of 1): TR / TE = 1000 / 33 ms, flip angle = 59°, resolution = 2.5 mm^3^ isotropic voxels, matrix size = 96 x 96, FoV = 240 x 240 mm, 48 axial slices with full cortical coverage and no gap, anterior-posterior phase encoding. At the beginning of each run, three dummy scans were acquired to allow for signal stabilization. At the beginning of each imaging session, a single dual-echo gradient-recalled echo (GRE) fieldmap was acquired for spatial distortion correction: TR = 675 ms, TE1/TE2 = 4.92/7.38 ms, flip angle = 60°, resolution = 2.5 mm^3^ isotropic voxels, matrix size = 96 x 96, 48 axial slices, anterior–posterior phase encoding. A T2-weighted structural scan was acquired with an in-plane acceleration factor of 4 using GRAPPA: TR/TE = 3200/563 ms, flip angle = 120°, resolution = 0.9375 x 0.9375 x 0.9 mm voxels, matrix size = 256 x 256, FoV = 240 x 240 x 172.8 mm, 192 sagittal slices, ascending acquisition, anterior–posterior phase encoding, no fat suppression, 3 min 21 s total acquisition time.

A T1-weighted scan acquired during the Grand Budapest Hotel session^17^ is included in this dataset. This scan had the following parameters: single-shot MPRAGE sequence with an in-plane acceleration factor of 2 using GRAPPA: TR / TE / TI = 2300 / 2.32 / 933 ms, flip angle = 8°, resolution = 0.9375 x 0.9375 x 0.9 mm voxels, matrix size = 256 x 256, FoV = 240 x 240 x 172.8 mm, 192 sagittal slices, ascending acquisition, anterior–posterior phase encoding, no fat suppression, 5 min 21 s total acquisition time.

Diffusion-weighted images were acquired using a single-shot spin-echo echo-planar imaging (SE-EPI) sequence with pre-scan normalization, fat suppression, and a multiband (i.e., simultaneous multi-slice; SMS) acceleration factor of 4: TR/TE = 2600/103 ms, flip angle = 90°, resolution = 2.0 mm^3^ isotropic voxels, matrix size = 120 x 120, FoV = 240 x 240 mm, 64 axial slices with no gap, posterior–anterior phase encoding. Diffusion weighting was applied along 96 non-collinear directions (HARDI scheme) with b = 1000 s/mm^2^, interleaved with 11 non-diffusion-weighted volumes (b = 0 s/mm^2^) distributed throughout the acquisition for a total of 107 volumes and approximately 4 min 38 s total acquisition time. For susceptibility-induced distortion correction of the diffusion-weighted images, a pair of spin-echo EPI scans was acquired with opposite phase encoding directions (anterior–posterior and posterior–anterior) using the same geometric parameters as the DWI acquisition: TR/TE = 9700/96 ms, flip angle = 90°, resolution = 2.0 mm^3^ isotropic voxels, matrix size = 120 x 120, FoV = 240 x 240 mm, 64 axial slices with no gap, fat suppression. Each phase encoding direction included 6 non-diffusion-weighted volumes (b = 0 s/mm^2^), with an effective echo spacing of 0.64 ms and total readout time of 76.2 ms.

### Dataset Preparation

DICOM data were converted to BIDS format (version 1.10.1) using HeuDiConv^35,36^ with the ReproIn heuristic.^37^ T1- and T2-weighted anatomical volumes were defaced using Pydeface.^38^

### Preprocessing

The text in this subsection is copied from the output of fMRIPrep^39,40^ as recommended by its developers (see https://www.nipreps.org/intro/transparency/#citation-boilerplates). Minor edits were performed for consistency and formatting.

Results included in this manuscript come from preprocessing performed using fMRIPrep 25.1.4^39,40^ (RRID:SCR_016216), which is based on Nipype 1.10.0^41,42^ (RRID:SCR_002502).

#### Preprocessing of B0 inhomogeneity mappings

A total of 2 fieldmaps were found available within the input BIDS structure for this particular subject. A B0 nonuniformity map (or fieldmap) was estimated from the phase-drift maps measure with two consecutive GRE (gradient-recalled echo) acquisitions. The corresponding phase-maps were phase-unwrapped with prelude (FSL).

#### Anatomical data preprocessing

A total of 1 T1-weighted (T1w) images were found within the input BIDS dataset. The T1w image was corrected for intensity non-uniformity (INU) with *N4BiasFieldCorrection*,^43^ distributed with ANTs 2.6.2^44^ (RRID:SCR_004757), and used as T1w-reference throughout the workflow. The T1w-reference was then skull-stripped with a Nipype implementation of the *antsBrainExtraction*.*sh* workflow (from ANTs), using OASIS30ANTs as target template. Brain tissue segmentation of cerebrospinal fluid (CSF), white-matter (WM) and gray-matter (GM) was performed on the brain-extracted T1w using *fast* (FSL, RRID:SCR_002823).^45^ Brain surfaces were reconstructed using *recon-all* (FreeSurfer 7.3.2, RRID:SCR_001847),^46^ and the brain mask estimated previously was refined with a custom variation of the method to reconcile ANTs-derived and FreeSurfer-derived segmentations of the cortical gray-matter of Mindboggle (RRID:SCR_002438).^47^ A T2-weighted image was used to improve pial surface refinement. Volume-based spatial normalization to one standard space (MNI152NLin2009cAsym) was performed through nonlinear registration with *antsRegistration* (ANTs 2.6.2), using brain-extracted versions of both T1w reference and the T1w template. The following template was selected for spatial normalization and accessed with TemplateFlow^48^ (24.2.2): ICBM 152 Nonlinear Asymmetrical template version 2009c^49^ (RRID:SCR_008796; TemplateFlow ID: MNI152NLin2009cAsym).

#### Functional data preprocessing

For each of the 16 BOLD runs found per subject (across all tasks and sessions), the following preprocessing was performed. First, a reference volume was generated, using a custom methodology of fMRIPrep, for use in head motion correction. Head-motion parameters with respect to the BOLD reference (transformation matrices, and six corresponding rotation and translation parameters) are estimated before any spatiotemporal filtering using *mcflirt* (FSL).^50^ The estimated fieldmap was then aligned with rigid-registration to the target EPI (echo-planar imaging) reference run. The field coefficients were mapped on to the reference EPI using the transform. The BOLD reference was then co-registered to the T1w reference using *bbregister* (FreeSurfer) which implements boundary-based registration.^51^ Co-registration was configured with six degrees of freedom. The aligned T2w image was used for initial co-registration. Several confounding time-series were calculated based on the preprocessed BOLD: framewise displacement (FD), DVARS and three region-wise global signals. FD was computed using two formulations following Power^52^ (absolute sum of relative motions) and Jenkinson^50^ (relative root mean square displacement between affines). FD and DVARS are calculated for each functional run, both using their implementations in Nipype (following the definitions by Power et al.^39^). The three global signals are extracted within the CSF, the WM, and the whole-brain masks. Additionally, a set of physiological regressors were extracted to allow for component-based noise correction (CompCor,^53^). Principal components are estimated after high-pass filtering the preprocessed BOLD time-series (using a discrete cosine filter with 128s cut-off) for the two CompCor variants: temporal (tCompCor) and anatomical (aCompCor). tCompCor components are then calculated from the top 2% variable voxels within the brain mask. For aCompCor, three probabilistic masks (CSF, WM and combined CSF+WM) are generated in anatomical space. The implementation differs from that of Behzadi et al. in that instead of eroding the masks by 2 pixels on BOLD space, a mask of pixels that likely contain a volume fraction of GM is subtracted from the aCompCor masks. This mask is obtained by dilating a GM mask extracted from the FreeSurfer’s aseg segmentation, and it ensures components are not extracted from voxels containing a minimal fraction of GM. Finally, these masks are resampled into BOLD space and binarized by thresholding at 0.99 (as in the original implementation). Components are also calculated separately within the WM and CSF masks. For each CompCor decomposition, the k components with the largest singular values are retained, such that the retained components’ time series are sufficient to explain 50 percent of variance across the nuisance mask (CSF, WM, combined, or temporal). The remaining components are dropped from consideration. The head-motion estimates calculated in the correction step were also placed within the corresponding confounds file. The confound time series derived from head motion estimates and global signals were expanded with the inclusion of temporal derivatives and quadratic terms for each ^54^. Frames that exceeded a threshold of 0.5 mm FD or 1.5 standardized DVARS were annotated as motion outliers. Additional nuisance timeseries are calculated by means of principal components analysis of the signal found within a thin band (crown) of voxels around the edge of the brain, as proposed by ref.^55^. The BOLD time-series were resampled onto the following surfaces (FreeSurfer reconstruction nomenclature): *fsaverage6*. All resamplings can be performed with a single interpolation step by composing all the pertinent transformations (i.e. head-motion transform matrices, susceptibility distortion correction when available, and co-registrations to anatomical and output spaces). Gridded (volumetric) resamplings were performed using *nitransforms*, configured with cubic B-spline interpolation. Non-gridded (surface) resamplings were performed using *mri_vol2surf* (FreeSurfer).

Many internal operations of fMRIPrep use Nilearn 0.11.1^56^ (RRID:SCR_001362), mostly within the functional processing workflow. For more details of the pipeline, see the section corresponding to workflows in fMRIPrep’s documentation.

### Data Validation

The functional data preprocessed by fMRIPrep were denoised using custom Python scripts. The following nuisance parameters were regressed out from the functional time series using ordinary least-squares regression: six motion parameters and their derivatives, global signal, framewise displacement, the first six noise components estimated by aCompCor, and polynomial trends up to second order. All metrics of interest were computed on data denoised as described, either in volume space or in surface space. No additional spatial smoothing or temporal filtering was performed. Temporal SNR, inter-subject correlation, and the other quality-control metrics were computed with these scripts using numpy,^57^ scipy,^58^ pandas,^59^ matplotlib,^60^ seaborn, nibabel,^61^ nilearn,^56^ scikit-learn,^62^ and pycortex.^63^

### Temporal Signal-to-Noise Ratio (tSNR)

We computed tSNR for each preprocessed functional run in native subject space (*space-T1w*) to avoid spatial smoothing from template normalization. For each voxel, tSNR was calculated as the temporal mean divided by the temporal standard deviation. To visualize tSNR variation across brain areas at the group level, we repeated the analysis on data resampled to the fsaverage6 surface.

### Inter-Subject Correlation (ISC)

ISC was computed to estimate the consistency of brain responses to the face clips. BOLD time series were projected to the template surface fsaverage6. Functional runs were reordered across participants to obtain the same stimulus order. Data corresponding to the initial buffer phase and memory test trials were discarded. To compute ISC, for each cortical vertex we correlated each participant’s time series with the mean time series of the remaining 20 participants. This procedure was repeated for all participants, and a median ISC map was computed at the group level.

## Data Records

To facilitate data sharing and the use of tools such as fMRIPrep and MRIQC,^39,66^ the unpreprocessed data were standardized following the Brain Imaging Data Structure^64^ (version 1.10.1). The hyperface raw dataset^67^ is available on OpenNeuro (https://openneuro.org/datasets/ds007329). This dataset includes the unpreprocessed data, as well as QA results and extensive derivative data used in ref.^18^ (see Figure 2). Additional fMRIPrep derivative data^68^ are available at https://openneuro.org/datasets/ds007384, and FreeSurfer derivative data^69^ are available at https://openneuro.org/datasets/ds007378. The raw data and derivatives are versioned and distributed with DataLad^70^ through OpenNeuro, and can be retrieved either directly from OpenNeuro or with DataLad. The fMRIPrep and FreeSurfer outputs are distributed as separate BIDS-Derivatives datasets. These derivatives are kept as separate, cross-linked datasets because the same participants also took part in the related studies (OpenNeuro ds003017,^65^ ds003834^9^), so some derivatives can be reused across these datasets without recomputation.

**Figure 2.**
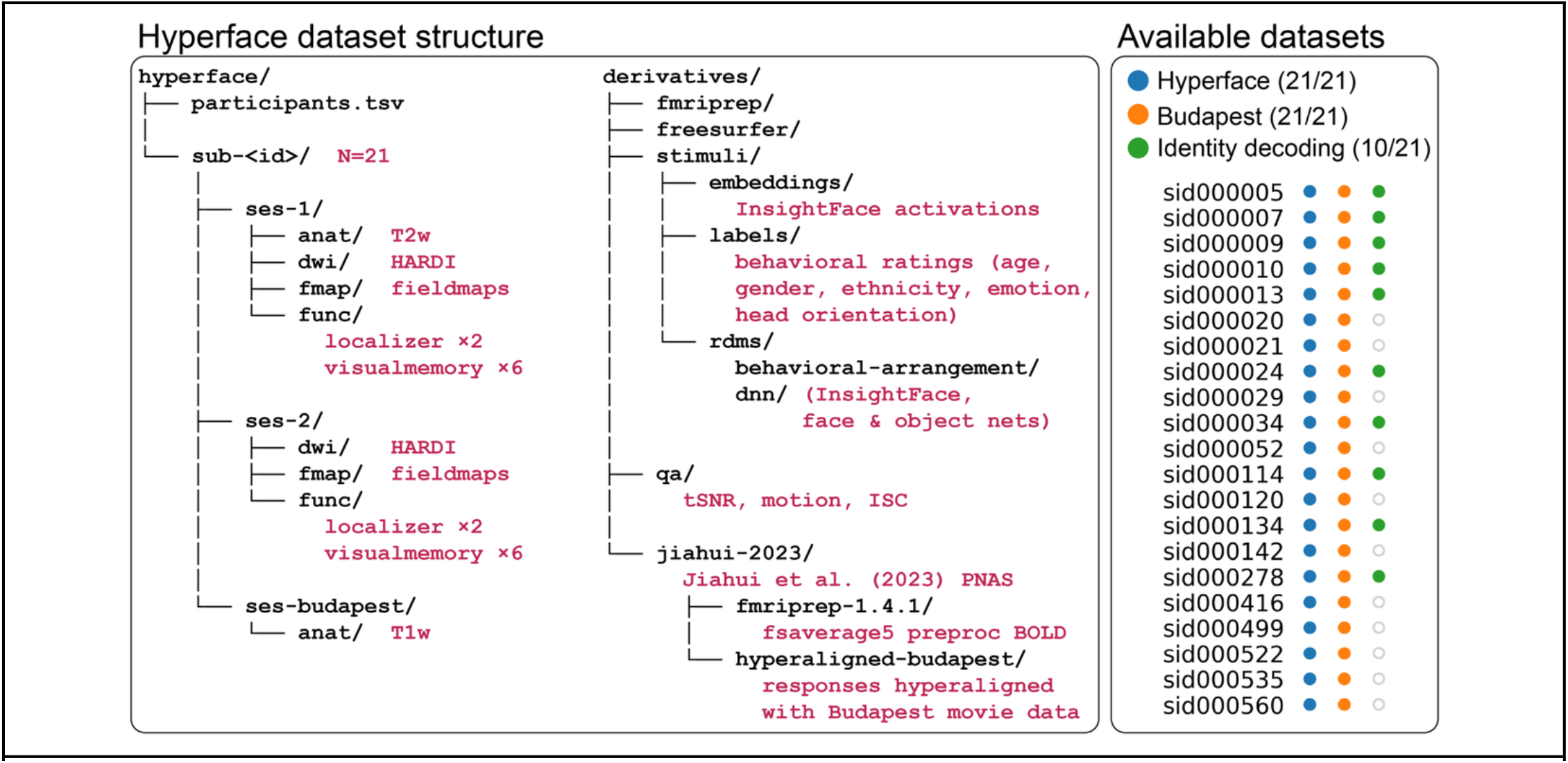
Dataset structure and available related datasets. **(Left)** The hyperface dataset follows the Brain Imaging Data Structure (BIDS) format.^64^ Raw data include two hyperface sessions (*ses-1, ses-2*), each comprising one GRE fieldmap scan, eight functional runs, and one diffusion-weighted imaging (DWI) scan; *ses-1* also includes a T2-weighted anatomical scan. A T1-weighted anatomical scan from a separate session (*ses-budapest*) is included from the publicly available Grand Budapest Hotel dataset.^17,65^ The derivatives folder contains two sets of preprocessed data: fMRIPrep 25.1.4 and FreeSurfer 7.3.2 outputs used for technical validation in this manuscript, and *jiahui-2023*, which includes preprocessed and hyperaligned data used in ref.^18^ Quality assurance (*qa*) files used for technical validation are also provided. Stimulus derivatives include InsightFace embeddings for all face clips, behavioral ratings from Mechanical Turk, and representational dissimilarity matrices (RDMs) derived from a behavioral arrangement task and from face- and object-specific deep neural networks (see ref.^18^ for details). **(Right)** Related publicly available datasets for the 21 hyperface participants. All participants also watched the second part of The Grand Budapest Hotel in the scanner.^17^ Ten participants completed two sessions of a controlled task used for face identity decoding with personally familiar and visually familiar faces.^9^ Together, these three fMRI datasets provide a unique resource for characterizing human face processing under naturalistic conditions and for benchmarking computer vision algorithms.

Presentation and QA scripts are available in a separate code repository^71^ at https://github.com/mvdoc/hyperface-data-paper. Because the face clips are excerpts of YouTube videos, the video files are not redistributed. Instead, the code repository provides a per-clip provenance manifest (source URL, in-video timing, and crop parameters) and code to regenerate and verify the stimuli.

## Technical Validation

We validated this dataset following our previously published approach to ensure data quality across separate domains.^17^ First, we analyzed participant motion to quantify behavioral sources of noise. Second, we estimated temporal signal-to-noise ratio (tSNR) for each voxel to verify comparable SNR levels across participants and to identify areas with low SNR. Third, we computed inter-subject correlation (ISC) as a check that the stimulus evoked consistent brain responses across participants.

### Motion

Participant motion was low throughout the dataset. The median framewise displacement across participants was 0.09 mm (range: 0.06–0.16 mm; Figure 3). The median percentage of volumes marked as motion outliers was 3.76% (range: 0.41–21.18%), with 13 of 21 participants having fewer than 5% outlier volumes. fMRIPrep defines an outlier as a volume with framewise displacement greater than 0.5 mm or standardized DVARS greater than 1.5 (see Methods).

**Figure 3.**
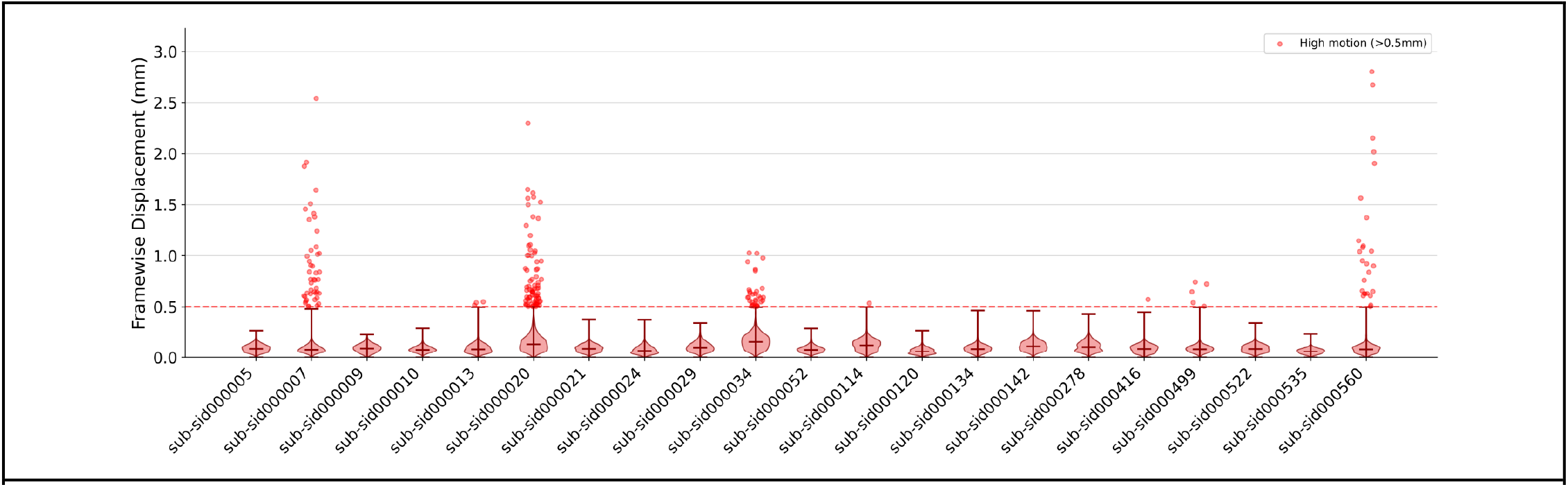
Framewise displacement for each participant across hyperface runs. Each violin plot shows the distribution of framewise displacement (FD) values across all functional volumes for one participant. Red dots indicate individual volumes with FD > 0.5 mm (dashed line). Subject motion was low across the dataset: median FD was 0.09 mm (range: 0.06–0.16 mm), well below the 0.5 mm threshold. Thirteen of 21 participants had fewer than 5% of volumes marked as motion outliers by fMRIPrep (FD > 0.5 mm or standardized DVARS > 1.5; see Methods).

### Temporal SNR

Temporal SNR (tSNR) was estimated for all participants in native anatomical space to minimize interpolation that would artificially inflate tSNR, and in *fsaverage6* template space for qualitative assessment across cortical areas. The mean whole-brain tSNR was 83.0 ± 4.0, comparable to previous datasets at 3T.^17,72,73^ As expected for a multiband BOLD EPI sequence, tSNR was highest in occipital and dorsal areas near the receiver coils and lowest in anterior temporal and orbitofrontal cortex near air-tissue boundaries (Figure 4).

**Figure 4.**
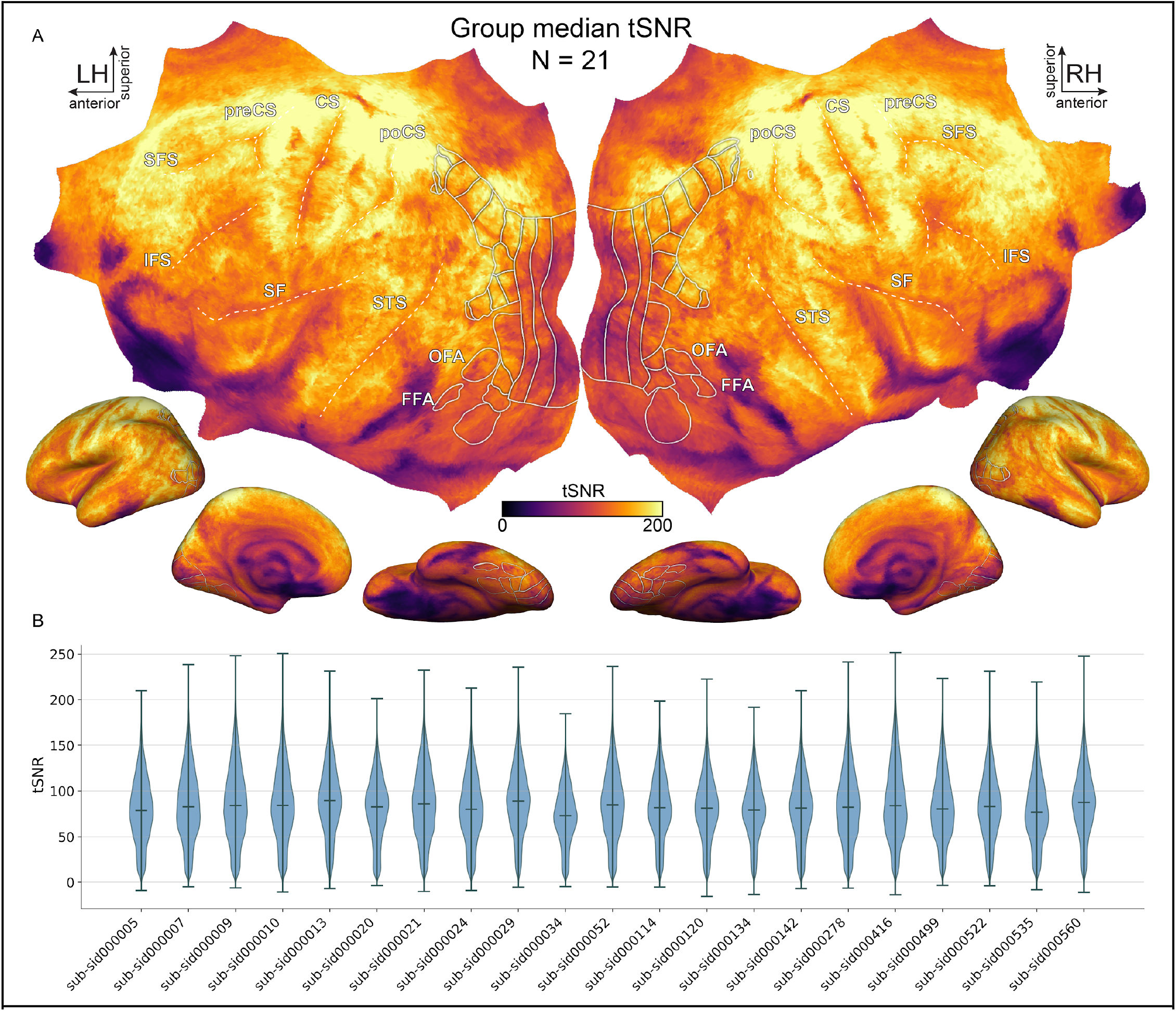
Temporal signal-to-noise ratio (tSNR) across participants. **A**. Group median tSNR (N=21) computed on data projected to *fsaverage6* surface space, displayed on a flatmap (top) and inflated surfaces (bottom). White outlines indicate anatomical landmarks and visual ROIs. tSNR is high across most of the cortex, with expected signal dropout near air-tissue boundaries in anterior temporal and orbitofrontal cortex. **B**. Violin plots show the distribution of voxelwise tSNR values for each participant, computed in native space. Group-averaged tSNR was 83.0 ± 4.0, consistent with other publicly available 3T datasets with similar voxel size. Abbreviations: CS, central sulcus; FFA, fusiform face area; IFS, inferior frontal sulcus; OFA, occipital face area; poCS, postcentral sulcus; preCS, precentral sulcus; SF, sylvian fissure; SFS, superior frontal sulcus; STS, superior temporal sulcus.

### Inter-Subject Correlation

ISC identifies regions where brain activity in response to the face clips was consistent across participants. Visual areas showed the highest ISC values, as expected from a dynamic visual stimulus (Figure 5). Positive ISC values were also found in face-processing regions such as the occipital face area (OFA), fusiform face area (FFA), and superior temporal sulcus (STS), as well as dorsal precuneus^8^ and right inferior frontal gyrus.^9,10^ Note that because this analysis was based on anatomical alignment, these ISC estimates represent a lower bound of what can be achieved with this dataset. Previous work has shown that hyperalignment^74–77^ provides much higher ISC values compared to anatomical alignment.^18,74^

**Figure 5.**
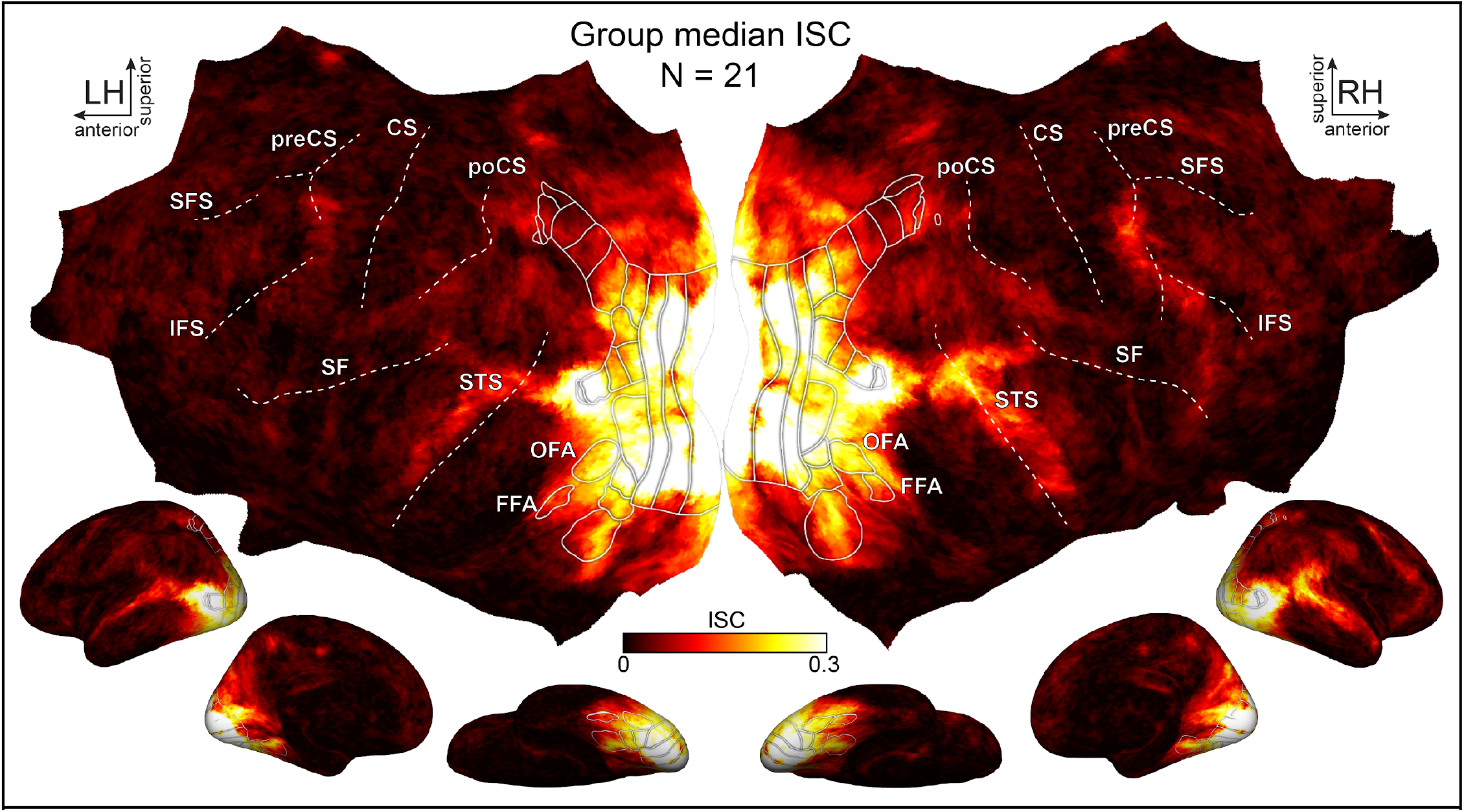
Group median inter-subject correlation (ISC). Flatmap and inflated surfaces show group median ISC (N=21) computed in *fsaverage6* space. White outlines indicate anatomical landmarks and visual ROIs. As expected from a dynamic visual stimulus, high ISC is found in primary visual and motion-sensitive areas. High ISC is also found in face-processing regions including the occipital face area (OFA), fusiform face area (FFA), and superior temporal sulcus (STS), as well as dorsal precuneus^8^ and right inferior frontal gyrus.^8–10^ Abbreviations: CS, central sulcus; FFA, fusiform face area; IFS, inferior frontal sulcus; OFA, occipital face area; poCS, postcentral sulcus; preCS, precentral sulcus; SF, sylvian fissure; SFS, superior frontal sulcus; STS, superior temporal sulcus.

Taken together, these analyses validate the high quality of this dataset. Low motion reflects good participant compliance during scanning. Comparable tSNR levels indicate consistent data quality across participants. ISC reveals shared brain responses in visual and face-processing areas, confirming that the naturalistic face stimuli elicited reliable brain activity across participants.

## Data Availability

The hyperface raw dataset is available on OpenNeuro (https://openneuro.org/datasets/ds007329). fMRIPrep and FreeSurfer derivatives are available at https://openneuro.org/datasets/ds007384 and https://openneuro.org/datasets/ds007378.

## Code Availability

All the code to replicate the QA analyses is available at https://github.com/mvdoc/hyperface-data-paper. The code uses several Python packages such as numpy,^57^ scipy,^58^ pandas,^59^ matplotlib,^60^ seaborn, nibabel,^61^ nilearn,^56^ scikit-learn,^62^ and pycortex.^63^ Additional code associated with our previous publication using this dataset^18^ is available at https://github.com/GUO-Jiahui/face_DCNN.

## Acknowledgments

We would like to thank Yaroslav Halchenko, Sam Nastase, Vassiki Chauhan, and the members of the Gobbini and Haxby labs for helpful discussions during the development of this project.

## Author Contributions

M.V.d.O.C. designed the experiment, wrote the presentation code, collected and analyzed the data, curated the data, and wrote the manuscript. G.J. and F.M. collected additional behavioral data, generated stimulus features, analyzed the data, and provided feedback on the manuscript. M.d.V. collected, edited, and curated the stimuli. M.I.G. and J.V.H. designed the experiment, obtained funding, and provided feedback on the manuscript. All authors read and approved the manuscript.

## References

1. Jack, R. E. & Schyns, P. G. The Human Face as a Dynamic Tool for Social Communication. Curr. Biol. 25, R621–R634 (2015).

2. Haxby, J. V., Hoffman, E. A. & Gobbini, M. I. Human neural systems for face recognition and social communication. Biol. Psychiatry 51, 59–67 (2002).

3. Haxby, J. V., Hoffman, E. A. & Gobbini, M. I. The distributed human neural system for face perception. Trends Cogn. Sci. 4, 223–233 (2000).

4. Todorov, A., Mandisodza, A. N., Goren, A. & Hall, C. C. Inferences of Competence from Faces Predict Election Outcomes. Science 308, 1623–1626 (2005).

5. Todorov, A., Gobbini, M. I., Evans, K. K. & Haxby, J. V. Spontaneous retrieval of affective person knowledge in face perception. Neuropsychologia 45, 163–173 (2007).

6. Kanwisher, N., McDermott, J. & Chun, M. M. The fusiform face area: a module in human extrastriate cortex specialized for face perception. J. Neurosci. 17, 4302–4311 (1997).

7. Haxby, J. V. et al. Distributed and overlapping representations of faces and objects in ventral temporal cortex. Science 293, 2425–2430 (2001).

8. Visconti di Oleggio Castello, M., Halchenko, Y. O., Swaroop Guntupalli, J., Gors, J. D. & Gobbini, M. I. The neural representation of personally familiar and unfamiliar faces in the distributed system for face perception. Sci. Rep. 7, 12237 (2017).

9. Visconti di Oleggio Castello, M., Haxby, J. V. & Gobbini, M. I. Shared neural codes for visual and semantic information about familiar faces in a common representational space. Proc. Natl. Acad. Sci. U. S. A. 118, e2110474118 (2021).

10. Guntupalli, J. S., Wheeler, K. G. & Gobbini, M. I. Disentangling the Representation of Identity from Head View Along the Human Face Processing Pathway. Cereb. Cortex 27, 46–53 (2017).

11. Duchaine, B. & Yovel, G. A Revised Neural Framework for Face Processing. Annual Review of Vision Science 1, 393–416 (2015).

12. Nastase, S. A., Goldstein, A. & Hasson, U. Keep it real: rethinking the primacy of experimental control in cognitive neuroscience. Neuroimage 222, 117254 (2020).

13. Matusz, P. J., Dikker, S., Huth, A. G. & Perrodin, C. Are We Ready for Real-world Neuroscience? J. Cogn. Neurosci. 31, 327–338 (2019).

14. Wu, M. C.-K., David, S. V. & Gallant, J. L. Complete functional characterization of sensory neurons by system identification. Annu. Rev. Neurosci. 29, 477–505 (2006).

15. Visconti di Oleggio Castello, M., Deniz, F., Dupré la Tour, T. & Gallant, J. L. Encoding models in functional magnetic resonance imaging: the Voxelwise Encoding Model framework. PsyArXiv (2025) doi:10.31234/osf.io/nt2jq_v2.

16. Chen, P. et al. An fMRI dataset in response to large-scale short natural dynamic facial expression videos. Sci Data 11, 1247 (2024).

17. Visconti di Oleggio Castello, M., Chauhan, V., Jiahui, G. & Gobbini, M. I. An fMRI dataset in response to ‘The Grand Budapest Hotel’, a socially-rich, naturalistic movie. Sci. Data 7, 383 (2020).

18. Jiahui, G. et al. Modeling naturalistic face processing in humans with deep convolutional neural networks. Proceedings of the National Academy of Sciences 120, e2304085120 (2023).

19. Tsantani, M. et al. FFA and OFA encode distinct types of face identity information. J. Neurosci. 41, 1952–1969 (2021).

20. Phillips, P. J. & White, D. The state of modelling face processing in humans with deep learning. Br. J. Psychol. 117, 656–676 (2026).

21. Visconti di Oleggio Castello, M. Characterizing Feature Representations in the Human Face-Processing Network with Multivariate Analyses and Encoding Models ([Unpublished doctoral …. (2018).

22. Nastase, S. A., Gazzola, V., Hasson, U. & Keysers, C. Measuring shared responses across subjects using intersubject correlation. Soc. Cogn. Affect. Neurosci. 14, 667–685 (2019).

23. Haxby, J. V., Connolly, A. C. & Guntupalli, J. S. Decoding neural representational spaces using multivariate pattern analysis. Annu. Rev. Neurosci. 37, 435–456 (2014).

24. Kriegeskorte, N., Mur, M. & Bandettini, P. Representational similarity analysis - connecting the branches of systems neuroscience. Front. Syst. Neurosci. 2, 4 (2008).

25. Dupré la Tour, T., Visconti di Oleggio Castello, M. & Gallant, J. L. The Voxelwise Encoding Model framework: A tutorial introduction to fitting encoding models to fMRI data. Imaging Neuroscience 3, (2025).

26. Naselaris, T., Kay, K. N., Nishimoto, S. & Gallant, J. L. Encoding and decoding in fMRI. Neuroimage 56, 400–410 (2011).

27. Ambadar, Z., Schooler, J. W. & Cohn, J. F. Deciphering the enigmatic face: the importance of facial dynamics in interpreting subtle facial expressions: The importance of facial dynamics in interpreting subtle facial expressions. Psychol. Sci. 16, 403–410 (2005).

28. Krumhuber, E. G., Kappas, A. & Manstead, A. S. R. Effects of dynamic aspects of facial expressions: A review. Emot. Rev. 5, 41–46 (2013).

29. O’Toole, A. J., Roark, D. A. & Abdi, H. Recognizing moving faces: a psychological and neural synthesis. Trends Cogn. Sci. 6, 261–266 (2002).

30. Kriegeskorte, N. & Mur, M. Inverse MDS: Inferring Dissimilarity Structure from Multiple Item Arrangements. Front Psychol 3, 245 (2012).

31. Nishimoto, S. et al. Reconstructing Visual Experiences from Brain Activity Evoked by Natural Movies. Curr. Biol. 21, 1641–1646 (2011).

32. Huth, A. G., Nishimoto, S., Vu, A. T. & Gallant, J. L. A continuous semantic space describes the representation of thousands of object and action categories across the human brain. Neuron 76, 1210–1224 (2012).

33. Nishimoto, S. & Gallant, J. L. A three-dimensional spatiotemporal receptive field model explains responses of area MT neurons to naturalistic movies. J. Neurosci. 31, 14551–14564 (2011).

34. Pitcher, D., Dilks, D. D., Saxe, R. R., Triantafyllou, C. & Kanwisher, N. Differential selectivity for dynamic versus static information in face-selective cortical regions. Neuroimage 56, 2356–2363 (2011).

35. Halchenko, Y. O. et al. HeuDiConv - flexible DICOM conversion into structured directory layouts. J Open Source Softw 9, (2024).

36. Halchenko, Y. O. et al. HeuDiConv — Flexible DICOM Conversion into Structured Directory Layouts. (Zenodo, 2025). doi:10.5281/ZENODO.17195198.

37. Visconti di Oleggio Castello, M. et al. ReproNim/Reproin: 0.11.6.2. (Zenodo, 2023). doi:10.5281/ZENODO.7975330.

38. Gulban, O. F. et al. Poldracklab/Pydeface: PyDeface v2.0.2. (Zenodo, 2022). doi:10.5281/ZENODO.6856482.

39. Esteban, O. et al. fMRIPrep: a robust preprocessing pipeline for functional MRI. Nat. Methods 16, 111–116 (2019).

40. Markiewicz, C. J., Esteban, O., Goncalves, M., Poldrack, R. A. & Gorgolewski, K. J. fMRIPrep: A Robust Preprocessing Pipeline for Functional MRI. (Zenodo, 2026). doi:10.5281/ZENODO.18248358.

41. Gorgolewski, K. et al. Nipype: a flexible, lightweight and extensible neuroimaging data processing framework in python. Front. Neuroinform. 5, 13 (2011).

42. Esteban, O. et al. Nipy/Nipype. (Zenodo, 2020). doi:10.5281/ZENODO.596855.

43. Tustison, N. J. et al. N4ITK: improved N3 bias correction. IEEE Trans. Med. Imaging 29, 1310–1320 (2010).

44. Avants, B. B., Epstein, C. L., Grossman, M. & Gee, J. C. Symmetric diffeomorphic image registration with cross-correlation: evaluating automated labeling of elderly and neurodegenerative brain. Med. Image Anal. 12, 26–41 (2008).

45. Zhang, Y., Brady, M. & Smith, S. Segmentation of brain MR images through a hidden Markov random field model and the expectation-maximization algorithm. IEEE Trans. Med. Imaging 20, 45–57 (2001).

46. Dale, A. M., Fischl, B. & Sereno, M. I. Cortical Surface-Based Analysis* 1:: I. Segmentation and Surface Reconstruction. Neuroimage 9, 179–194 (1999).

47. Klein, A. et al. Mindboggling morphometry of human brains. PLoS Comput. Biol. 13, e1005350 (2017).

48. Ciric, R. et al. TemplateFlow: FAIR-sharing of multi-scale, multi-species brain models. Nat Methods 19, 1568–1571 (2022).

49. Fonov, V. S., Evans, A. C., McKinstry, R. C., Almli, C. R. & Collins, D. L. Unbiased nonlinear average age-appropriate brain templates from birth to adulthood. Neuroimage Supplement 1, S102 (2009).

50. Jenkinson, M., Bannister, P., Brady, M. & Smith, S. Improved optimization for the robust and accurate linear registration and motion correction of brain images. Neuroimage 17, 825–841 (2002).

51. Greve, D. N. & Fischl, B. Accurate and robust brain image alignment using boundary-based registration. Neuroimage 48, 63–72 (2009).

52. Power, J. D. et al. Methods to detect, characterize, and remove motion artifact in resting state fMRI. Neuroimage 84, 320–341 (2014).

53. Behzadi, Y., Restom, K., Liau, J. & Liu, T. T. A component based noise correction method (CompCor) for BOLD and perfusion based fMRI. Neuroimage 37, 90–101 (2007).

54. Satterthwaite, T. D. et al. An improved framework for confound regression and filtering for control of motion artifact in the preprocessing of resting-state functional connectivity data. Neuroimage 64, 240–256 (2013).

55. Patriat, R., Reynolds, R. C. & Birn, R. M. An improved model of motion-related signal changes in fMRI. Neuroimage 144, 74–82 (2017).

56. Abraham, A. et al. Machine learning for neuroimaging with scikit-learn. Front. Neuroinform. 8, 14 (2014).

57. van der Walt, S., Colbert, S. C. & Varoquaux, G. The NumPy Array: A Structure for Efficient Numerical Computation. Comput. Sci. Eng. 13, 22–30 (2011).

58. Virtanen, P. et al. SciPy 1.0: fundamental algorithms for scientific computing in Python. Nat. Methods 17, 261–272 (2020).

59. McKinney, W. Data Structures for Statistical Computing in Python. in Proceedings of the 9th Python in Science Conference (eds van der Walt, S. & Millman, J.) 56–61 (2010).

60. Hunter, J. D. Matplotlib: A 2D Graphics Environment. Comput. Sci. Eng. 9, 90–95 (2007).

61. Brett, M. et al. Nipy/Nibabel. (Zenodo, 2024). doi:10.5281/zenodo.591597.

62. Pedregosa, F. et al. Scikit-learn: Machine Learning in Python. J. Mach. Learn. Res. abs/1201.0490, (2011).

63. Gao, J. S., Huth, A. G., Lescroart, M. D. & Gallant, J. L. Pycortex: an interactive surface visualizer for fMRI. Front. Neuroinform. 9, 23 (2015).

64. Gorgolewski, K. J. et al. The brain imaging data structure, a format for organizing and describing outputs of neuroimaging experiments. Scientific Data 3, 160044 (2016).

65. Visconti di Oleggio Castello, M., Chauhan, V., Jiahui, G. & Gobbini, M. I. Budapest. OpenNeuro 10.18112/openneuro.ds003017.v1.0.0 (2020).

66. Esteban, O. et al. MRIQC: Advancing the automatic prediction of image quality in MRI from unseen sites. PLoS One 12, e0184661 (2017).

67. Visconti di Oleggio Castello, M. et al. hyperface: a naturalistic fMRI dataset to characterize human face processing. OpenNeuro 10.18112/openneuro.ds007329.v1.0.2 (2026).

68. Visconti di Oleggio Castello, M. et al. fMRIPrep derivatives for hyperface: a naturalistic fMRI dataset to characterize human face processing. OpenNeuro 10.18112/openneuro.ds007384.v1.0.0 (2026).

69. Visconti di Oleggio Castello, M. et al. FreeSurfer derivatives for hyperface: a naturalistic fMRI dataset to characterize human face processing. OpenNeuro 10.18112/openneuro.ds007378.v1.0.0 (2026).

70. Halchenko, Y. O. et al. DataLad: distributed system for joint management of code, data, and their relationship. J. Open Source Softw. 6, 3262 (2021).

71. Visconti di Oleggio Castello, M. et al. Code for Hyperface: A Naturalistic fMRI Dataset for Investigating Human Face Processing. (Zenodo, 2026). doi:10.5281/zenodo.21188776.

72. Nastase, S. A. et al. The “Narratives” fMRI dataset for evaluating models of naturalistic language comprehension. Scientific Data 8, 1–22 (2021).

73. Aliko, S., Huang, J., Gheorghiu, F., Meliss, S. & Skipper, J. I. A naturalistic neuroimaging database for understanding the brain using ecological stimuli. Sci Data 7, 347 (2020).

74. Haxby, J. V., Guntupalli, J. S., Nastase, S. A. & Feilong, M. Hyperalignment: Modeling shared information encoded in idiosyncratic cortical topographies. Elife 9, (2020).

75. Haxby, J. V. et al. A Common, High-Dimensional Model of the Representational Space in Human Ventral Temporal Cortex. Neuron 72, 404–416 (2011).

76. Guntupalli, J. S. et al. A model of representational spaces in human cortex. Cereb. Cortex 26, 2919–2934 (2016).

77. Feilong, M. et al. The Individualized Neural Tuning Model: Precise and generalizable cartography of functional architecture in individual brains. Imaging Neuroscience (2023) doi:10.1162/imag_a_00032.

